# An Atlas of Brain–Bone Sympathetic Neural Circuits

**DOI:** 10.1101/2024.02.07.579382

**Authors:** Vitaly Ryu, Anisa Gumerova, Ronit Witztum, Funda Korkmaz, Hasni Kannangara, Ofer Moldavski, Orly Barak, Daria Lizneva, Ki A. Goosens, Sarah Stanley, Se-Min Kim, Tony Yuen, Mone Zaidi

**Affiliations:** Center for Translational Medicine and Pharmacology (CeTMaP), Icahn School of Medicine at Mount Sinai, New York, NY 10029; Department of Medicine and of Pharmacological Sciences, Icahn School of Medicine at Mount Sinai, New York, NY 10029; Department of Psychiatry, Icahn School of Medicine at Mount Sinai, New York, NY 10029

## Abstract

There is clear evidence that the sympathetic nervous system (SNS) mediates bone metabolism. Histological studies show abundant SNS innervation of the periosteum and bone marrow––these nerves consist of noradrenergic fibers that immunostain for tyrosine hydroxylase, dopamine beta hydroxylase, or neuropeptide Y. Nonetheless, the brain sites that send efferent SNS outflow to bone have not yet been characterized. Using pseudorabies (PRV) viral transneuronal tracing, we report, for the first time, the identification of central SNS outflow sites that innervate bone. We find that the central SNS outflow to bone originates from 87 brain nuclei, sub–nuclei and regions of six brain divisions, namely the midbrain and pons, hypothalamus, hindbrain medulla, forebrain, cerebral cortex, and thalamus. We also find that certain sites, such as the raphe magnus (RMg) of the medulla and periaqueductal gray (PAG) of the midbrain, display greater degrees of PRV152 infection, suggesting that there is considerable site–specific variation in the levels of central SNS outflow to bone. This comprehensive compendium illustrating the central coding and control of SNS efferent signals to bone should allow for a greater understanding of the neural regulation of bone metabolism, and importantly and of clinical relevance, mechanisms for central bone pain.

## INTRODUCTION

Elegant studies have suggested that increased sympathetic nervous system (SNS) tone causes bone loss through a reduction in bone formation, which is coupled with increased bone resorption (Elefteriou, 2018; Elefteriou *et al*, 2005; Takeda *et al*, 2002). It has also been shown using leptin–deficient mice with a high bone mass that the anti–osteogenic actions of leptin are mediated centrally by glucose–responsive neurons in the ventromedial hypothalamus through peripheral SNS pathways (Ducy *et al*, 2000; Takeda *et al*., 2002). Furthermore, both the periosteum and the bone marrow are innervated richly by the SNS as evidenced by immunoreactive tyrosine hydroxylase, dopamine beta hydroxylase, or neuropeptide Y fibers. These latter SNS markers are associated mostly with the vasculature and SNS vesicular acetylcholine transporter (VAChT), whereas vasoactive intestinal polypeptide (VIP) immunoreactive fibers display mainly a parenchymal location (Francis *et al*, 1997; Hill & Elde, 1991; Hohmann *et al*, 1986; Martin *et al*, 2007). Despite these studies, the distribution of SNS nerves within the mammalian skeleton and their connectivity to central neurons is far from being completely understood.

Viral transneuronal tracing has become an established technology to define central SNS outflow circuitry to peripheral organs. Bartha’s K strain of the pseudorabies virus (PRV) is a transneuronal tract tracer that provides the ability to map multi–synaptic circuits within the same animal (Ekstrand *et al*, 2008; Enquist, 2002; Song *et al*, 2005a). Once in the host, PRVs are endocytosed at axon terminal membranes after binding to viral attachment proteins, which act as ‘viral receptors’. Transported exclusively in a retrograde manner from the dendrites of the infected neurons to axons, PRVs first make synaptic contact with neuronal cell bodies and undergo self–amplification and thereafter continue their specific backward ascent (Curanovic & Enquist, 2009). This results in an infection that progresses along the neuroaxis chain from the periphery to higher CNS sites (Ekstrand *et al*., 2008; Enquist, 2002; Song *et al*., 2005a).

Utilizing this viral technology, we have previously shown postganglionic SNS innervation of specific white and brown adipose tissue depots with the separate and shared central SNS relay sites (Ryu & Bartness, 2014; Ryu *et al*, 2015). Moreover, we have established a direct neuroanatomical connection between phosphodiesterase 5A (PDE5A)–containing neurons in specific brain nuclei and bone, inferring a contribution of the central nodes to the bone–forming actions of PDE5A inhibitors (Kim *et al*, 2020). A hierarchical circuit controlling SNS output to rat femoral epiphyseal bone marrow has also been defined by identifying PRVs in ganglia and paravertebral chain in the intermediolateral column of the lower thoracic spinal cord (Denes *et al*, 2005). In addition, neurons in C1, A5, A7 catecholaminergic cell groups and several other nuclei of the ventrolateral and ventromedial medulla, periaqueductal gray, the paraventricular hypothalamic nucleus, among other hypothalamic nuclei, as well as the insular and piriform cortex comprise the known central network sending SNS outflow to bone marrow (Denes *et al*., 2005). However, no studies have yet mapped the exact localization and organization of the central SNS circuitry innervating the murine femur. The purpose of the present study was thus to identify central SNS sites innervating bone and to investigate whether separate or/and shared central SNS circuitries underpin the autonomic mediation of bone.

## METHODS

### Mice

Adult male mice (∼3 to 4–month–old) were single–housed in a 12 h:12 h light:dark cycle at 22 ± 2 ºC with *ad libitum* access to water and regular chow. All procedures were approved by the Mount Sinai Institutional Animal Care and Use Committee and were performed in accordance with Public Health Service and United States Department of Agriculture guidelines.

### Viral injections

To identify brain sites sending the SNS outflow to bone, we used a transsynaptic tracing technique with a pseudorabies virus strain, PRV152. PRV152 expresses enhanced green fluorescent protein (EGFP) under control of the human cytomegalovirus immediate–early promoter. When injected into peripheral tissues, the virus travels exclusively in a retrograde manner and localizes to central neurons, thus allowing the mapping of the entire periphery– brain neuroaxis.

All virus injections were performed according to Biosafety Level 2 standards. Mice (*N*=6) were anesthetized with isoflurane (2–3% in oxygen; Baxter Healthcare, Deerfield, IL) and the right femur–tibia joint was exposed for a series of PRV152 microinjections (4.7 × 10^9^ pfu/mL) into five loci (150 nL/locus) evenly distributed across the bone metaphysis and periosteum areas, which are known to be enriched with SNS innervation. The syringe was held in place for 60 seconds to prevent efflux of virus after each injection. Finally, the incision was closed with sterile sutures and wound clips. Nitrofurozone powder (nfz Puffer; Hess & Clark, Lexington, KY) was applied locally to minimize the risk of bacterial infection. Note that, as a control for viral injection, we showed that no EGFP signal was detected when PRV152 was placed on the bone surface rather than injected into the periosteum or metaphyseal bone. In addition, we found PRV152–infected neurons in the intermediolateral cell column (IML) of the spinal cords, suggesting specific bone–SNS ganglia–IML–brain route of infection, which is in concordance with our previous findings where PRV152 individually infected the classic SNS spinal cord neurons (Bamshad *et al*, 1999; Ryu & Bartness, 2014; Ryu *et al*., 2015).

### Histology

Animals were sacrificed 6 days after the last PRV152 injection based on the progression of both viruses to the brain in pilot studies (Ryu, V., unpublished observations). Mice were euthanized with carbon dioxide and perfused transcardially with 0.9% heparinized saline followed by 4% paraformaldehyde in 0.1 M phosphate–buffered saline (PBS; pH 7.4). Brains were collected and post–fixed in the same fixative for 3 to 4 hours at 4 °C, then transferred to a 30% sucrose solution in 0.1 M PBS with 0.1 % sodium azide and stored at 4 °C until sectioning on a freezing stage sliding microtome at 25 μm. Sections were stored in 0.1 M PBS solution with 0.1% sodium azide until processing for immunofluorescence.

For immunofluorescence, free–floating brain sections were rinsed in 0.1 M PBS (2 × 15 minutes) followed by a 30-minute blocking in 10% normal goat serum (NGS; Vector Laboratories, Burlingame, CA) and 0.4% Triton X-100 in 0.1 M PBS. Next, sections were incubated with a primary chicken anti–EGFP antibody (1:1000; Thermo Fisher Scientific, catalog no. A10262) for 18 hours. Sections were then incubated in the secondary AlexaFluor-488-coupled goat anti–chicken antibody (1:700; Jackson Immunoresearch, catalog no. 103-545-155) with 2% NGS and 0.4% Triton X-100 in 0.1 M PBS at room temperature for 2 hours. For immunofluorescence controls, the primary antibody was either omitted or pre–adsorbed with the immunizing peptide overnight at 4 °C resulting in no immunoreactive staining. Sections were mounted onto slides (Superfrost Plus) and cover–slipped using ProLong Gold Antifade Reagent (Thermo Fisher Scientific, catalog no. P36982). All steps were performed at room temperature.

### Quantitation

Immunofluorescence images were viewed and captured using x10 and x20 magnification with an Observer.Z1 fluorescence microscope (Carl Zeiss, Germany) with appropriate filters for AlexaFluor-488 and DAPI. The single–labeled PRV152 and DAPI images were evaluated and overlaid using Zen software (Carl Zeiss, Germany) and ImageJ (NIH, Bethesda, MD). We counted cells positive for SNS PRV152 immunoreactivity in every sixth brain section using the manual tag feature of the Adobe Photoshop CS5.1 software, thus eliminating the likelihood of counting the same neurons more than once. Neuron numbers in the brain were averaged across each examined nucleus/sub–nucleus/region from all animals. A mouse brain atlas (Paxinos and Franklin, 2007) was used to identify brain areas. For the photomicrographs, we used Adobe Photoshop CS5.1 (Adobe Systems) only to adjust the brightness, contrast and sharpness, to remove artifactual obstacles (i.e., obscuring bubbles) and to make the composite plates.

## RESULTS

### Validation

Following PRV152 infections, mice remained asymptomatic until day 5 post–inoculation, after which time, mice began to display symptoms of infection, including occasional loss of body weight and decreased mobility, but most often an ungroomed coat. Mice were euthanized for histological analyses when such symptoms became apparent. Four of six mice were equally infected by PRV152 throughout the neuroaxis from the hindbrain to the forebrain and therefore were included in the analyses. Two mice exhibited over–infection by PRV152, as evidenced by widespread cloudy plaques surrounding the infected neurons; these mice were excluded from the analysis. We also found PRV152–labeled neurons in the IML of the spinal cord in accordance with our previous studies that defined SNS innervations of fat pads in the Siberian hamster (Ryu & Bartness, 2014; Ryu *et al*., 2015; Ryu *et al*, 2017).

Unilateral PRV152 microinjection into the right femur appeared bilaterally in the brain with almost no noticeable domination of the viral infection between the two hemispheres. Likewise, prior studies on SNS and sensory innervations of various fat depots, utilizing the SNS tract tracer PRV152 and sensory system tract tracer HSV-1 produced no ipsilateral differences between the innervation patterns of SNS or sensory system with unilateral viral inoculation (Bamshad *et al*., 1999; Leitner & Bartness, 2009; Song *et al*, 2008; Vaughan & Bartness, 2012).

To validate the retrograde tract tracing methodology, we placed PRV152 at the same titer on the bone surface, rather than injecting it into the periosteum or metaphysis. No EGFP signal was detected in the PVH that is known to possess sympathetic pre–autonomic neurons or in the RPa (Fig. 1A). By contrast, PRV152 injections into the periosteum or metaphysis resulted in positive EGFP immunostaining in the PVH (Fig. 1B). In addition, we found PRV152– infected neurons in the IML of the spinal cord, at T13–L2 levels (Fig. 1B), suggesting specific bone–SNS ganglia–IML–brain route of infection; this is consistent with prior findings wherein PRV152 individually infected the classic SNS spinal cord neurons (Bamshad *et al*., 1999; Ryu & Bartness, 2014; Ryu *et al*., 2015).

**Figure 1:**
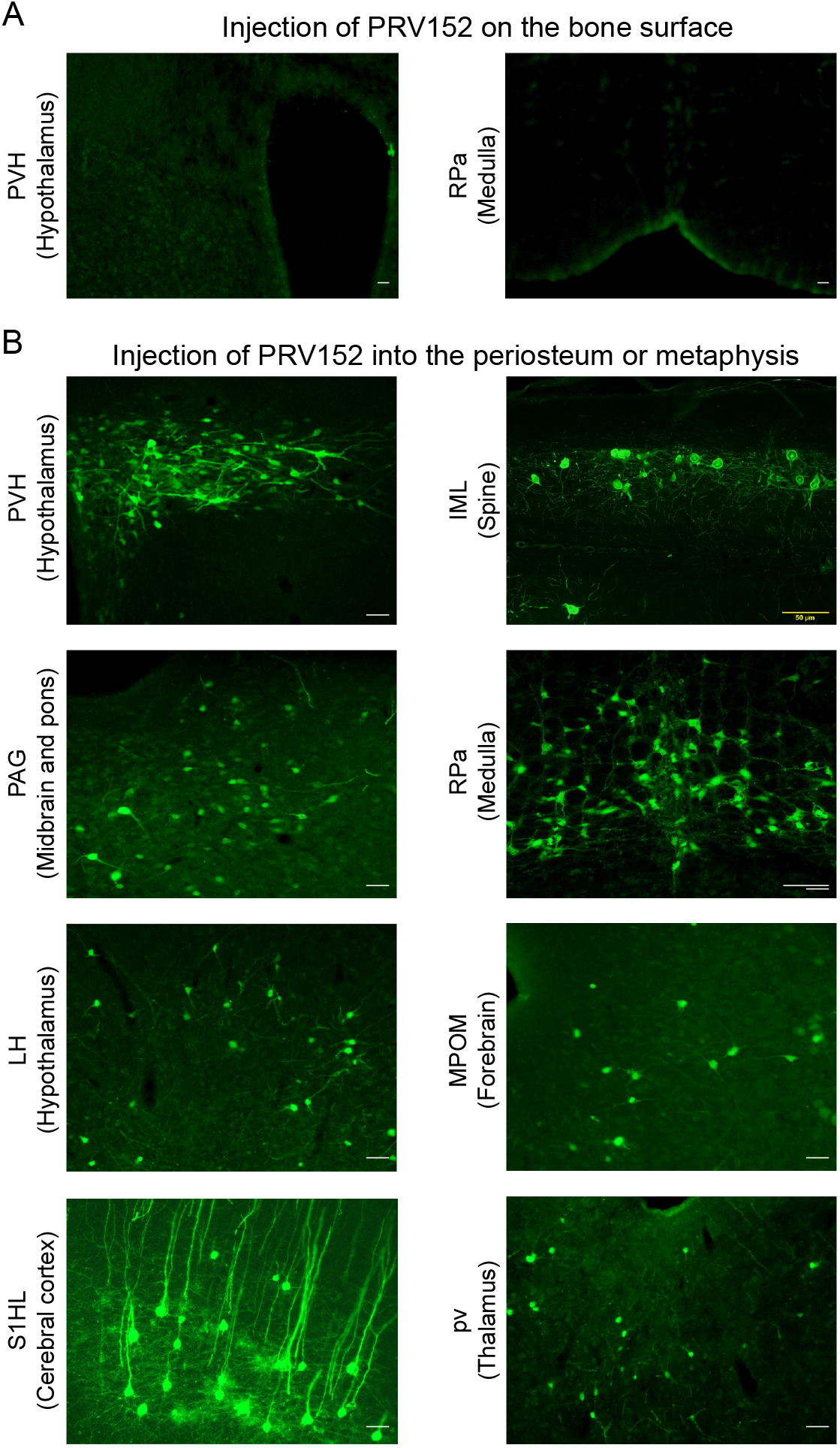
PRV152 transneuronal viral tract tracing. As a control for viral injection, no EGFP signal was detected in the PVH, known to possess main sympathetic pre-autonomic neurons, and the RPa, when PRV152 was placed on the bone surface. By contrast, PRV152 injections into the periosteum or metaphyseal bone resulted in positive EGFP immunoreactivity in the PVH. In addition, we found PRV152–infected neurons in the intermediolateral cell column (IML) of the spinal cord at T13–L2 levels, suggesting specific bone–SNS ganglia–IML–brain route of infection which are in concordance with our previous findings where PRV152 individually infected the classic SNS spinal cord neurons. Also shown are representative microphotograph illustrating PRV152 immunolabeling in the PAG (midbrain and pons), RPa (medulla), LH (hypothalamus), MPOM (forebrain), S1HL (cerebral cortex), and pv (thalamus). PVH, paraventricular hypothalamic nucleus; PAG, periaqueductal gray; RPa, raphe pallidus; LH, lateral hypothalamus; MPOM, medial preoptic nucleus, medial part; S1HL, primary somatosensory cortex, hindlimb region; pv, periventricular fiber system. Scale bar = 50 µm.

### Viral Infections in The Brain

We identified 87 PRV152–positive brain nuclei, sub–nuclei and regions within six brain divisions, with the hypothalamus having the most PRV152–infected SNS neurons connecting to bone (1177.25 ± 62.75), followed, in descending order, by midbrain and pons (1065 ± 22.39), hindbrain medulla (495.25 ± 33.49), forebrain (237.5 ± 15.08), cerebral cortex (104.75 ± 4.64) and thalamus (65.25 ± 7.78) (Fig. 2). Hypothalamic areas with the highest percentages of PRV152–labeled neurons included the lateral hypothalamus (LH), PVH and dorsomedial hypothalamus (DM) (Fig. 1B and Fig. 2; see Appendix for a glossary of brain nuclei, sub-nuclei and regions). The LH and PVH also were among the regions with the highest absolute numbers of infected neurons of the 25 PRV152–positive nuclei, sub–nuclei, and regions. In the midbrain and pons, areas with the highest percentages and counts of PRV152–infected neurons included the PAG, lateral PAG (LPAG) and pontine reticular nucleus, oral part (PnO), among 18 nuclei, sub–nuclei, and regions. Single–labeled neurons were also notable in the hindbrain medulla, where the raphe pallidus nucleus (RPa), RMg, and gigantocellular reticular nucleus (Gi) were among 23 nuclei, sub–nuclei and regions, heavily represented by the largest percentages and counts of PRV152–labeled neurons. The forebrain areas with the highest percentages and numbers of PRV152–labeled neurons were the medial preoptic nucleus, medial part (MPOM), bed nucleus of the stria terminalis (BST) and lateral septal nucleus, ventral part (LSV) among 15 PRV152–positive nuclei, sub–nuclei, and regions. In the cerebral cortex, there were only 3 regions containing PRV152–labeled neurons––namely, the primary somatosensory cortex, hindlimb region (S1HL), secondary and primary motor cortex (M2 and M1, respectively). The S1HL and M2 had both the highest percentages and numbers of PRV152–labeled neurons. Finally, we detected 3 brain sites with PRV152–infected neurons within the thalamus. Among the nuclei possessing the highest percentages and numbers of PRV152–labeled neurons were the periventricular fiber system (pv) and precommisural nucleus (PrC).

**Figure 2:**
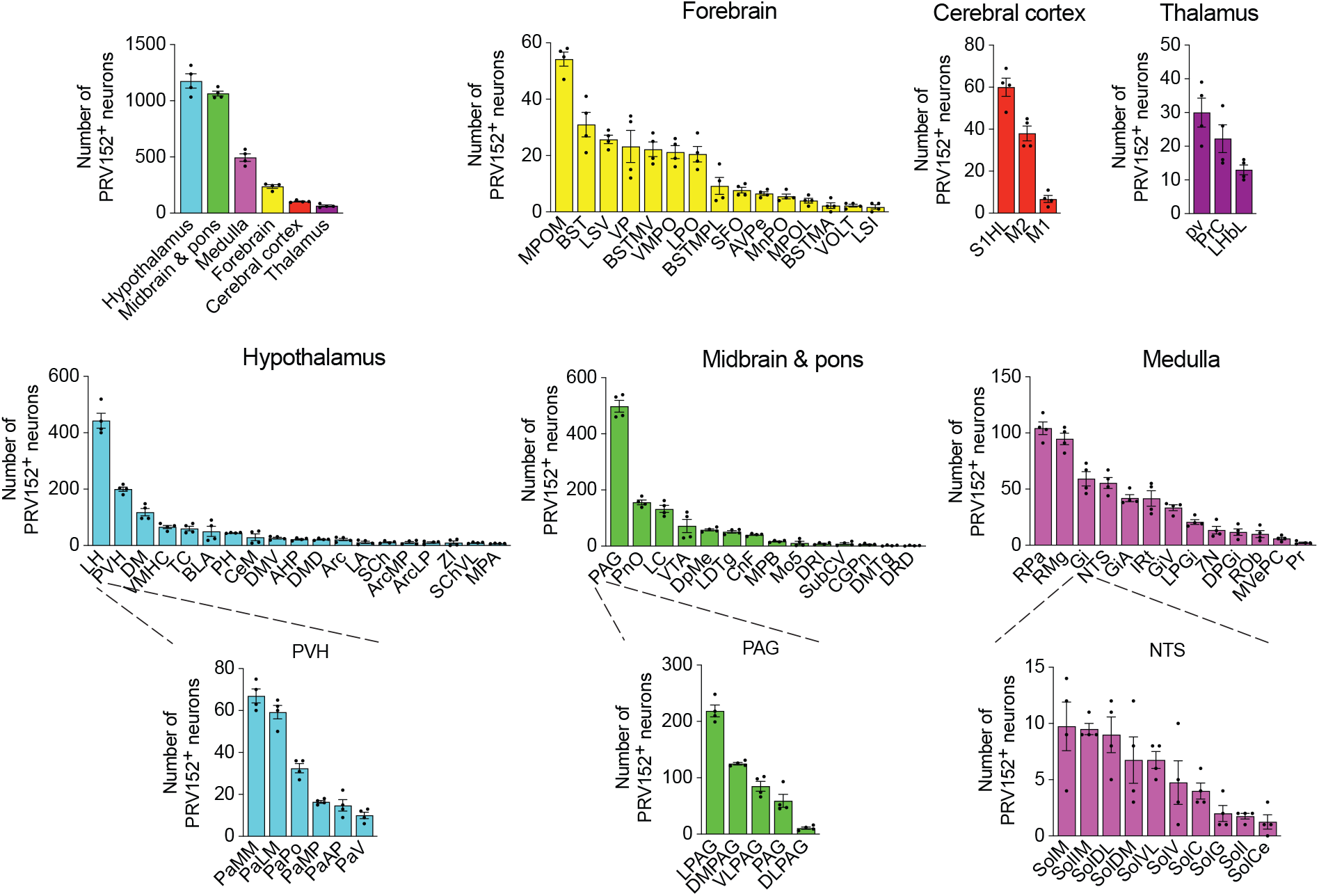
PRV152 immunolabeling in brain regions, sub-regions and nuclei. Numbers of PRV152–labeled neurons in brain regions, namely, hypothalamus, midbrain and pons, medulla, forebrain, cerebral cortex and thalamus, as well as their sub-regions and nuclei, following viral injections into bone.

## DISCUSSION

Using transneuronal tract tracers, we (Ryu & Bartness, 2014; Ryu *et al*., 2015; Ryu *et al*., 2017) and others (Bamshad *et al*, 1998; Bowers *et al*, 2004; Shi & Bartness, 2001; Song & Bartness, 2001) have documented postganglionic SNS innervation of white and brown adipose tissue depots with the separate and shared central SNS nodes. Moreover, we have recently established a direct neuroanatomical link between PDE5A–containing neurons in specific brain sites and bone (Kim *et al*., 2020). We report here, for the first time, a comprehensive atlas that defines with remarkable precision the crosstalk between the SNS and bone. Notably, the PRV152 neural tract tracer especially predominate in the PAG of the midbrain, LH of the hypothalamus, RPa of the medulla, MPOM of the forebrain, S1HL of the cortex and pv of the thalamus. Collectively, these data provide important insights into the distributed neural system integrating SNS neural circuitry with bone.

Neuroanatomical and functional evidence in mice suggests that the SNS regulates bone remodeling and bone mass (Ducy *et al*., 2000; Francis *et al*., 1997; Hill & Elde, 1991; Hohmann *et al*., 1986; Martin *et al*., 2007; Takeda *et al*., 2002). Furthermore, it is clear that leptin acts as an anti–osteogenesis signal through glucose responsive neurons in the VMH *via* peripheral SNS relay (Takeda *et al*., 2002). These data are consistent with histological evidence, using SNS markers in noradrenergic fibers, for a rich innervation of the periosteum and of bone marrow (Francis *et al*., 1997; Hill & Elde, 1991; Hohmann *et al*., 1986; Martin *et al*., 2007). Likewise, dopamine–transporter–deficient mice with no rapid uptake of dopamine into presynaptic terminals are osteopenic (Bliziotes *et al*, 2000). Multisynaptic tract tracing has identified limited hierarchical central circuitry controlling SNS innervation of rat femoral epiphyseal bone marrow and bone (Denes *et al*., 2005). Several SNS pathways from the brainstem and the hypothalamus relay to femoral bone marrow and the femur through preganglionic neurons in the lower thoracic and upper lumbar segments T4 to L1 of the IML and postganglionic neurons in paravertebral chain ganglia at lumbar levels (Denes *et al*., 2005).

Despite the fact that largely the same brain sites project to both the femur (our findings) and bone marrow (Denes *et al*., 2005), some sites display higher levels of PRV152 infectivity than others––this suggests that separate site–specific SNS circuits may project to the femur and femoral bone marrow. These overlapping SNS–innervating circuits to both sites include the midbrain PAG, somatosensory cortex, forebrain MPOM, thalamic periventricular nucleus, hypothalamic PVH and lateral hypothalamic nucleus (LA), and medulla RPa. The PAG receives afferent fibers not only from the parabrachial nucleus and RPa (Mantyh, 1982), which contain PDE5A–expressing neurons sending SNS outputs to bone (Kim *et al*., 2020), but also from the spinal cord (Pechura & Liu, 1986). We and others have previously shown that the PAG sends SNS outflow to WAT in Siberian hamsters (Bamshad *et al*., 1998; Nguyen *et al*, 2014; Ryu & Bartness, 2014; Song *et al*, 2005b) and the laboratory rat (Adler *et al*, 2012). Most notably, the PAG is largely responsible for SNS responses and descending modulation of pain perception (Baptista-de-Souza *et al*, 2018; Benarroch, 2008; Calvino & Grilo, 2006). Therefore, this midbrain node could receive sensory inflow relating to bone pain and provide SNS relay (Fig. 3).

**Figure 3:**
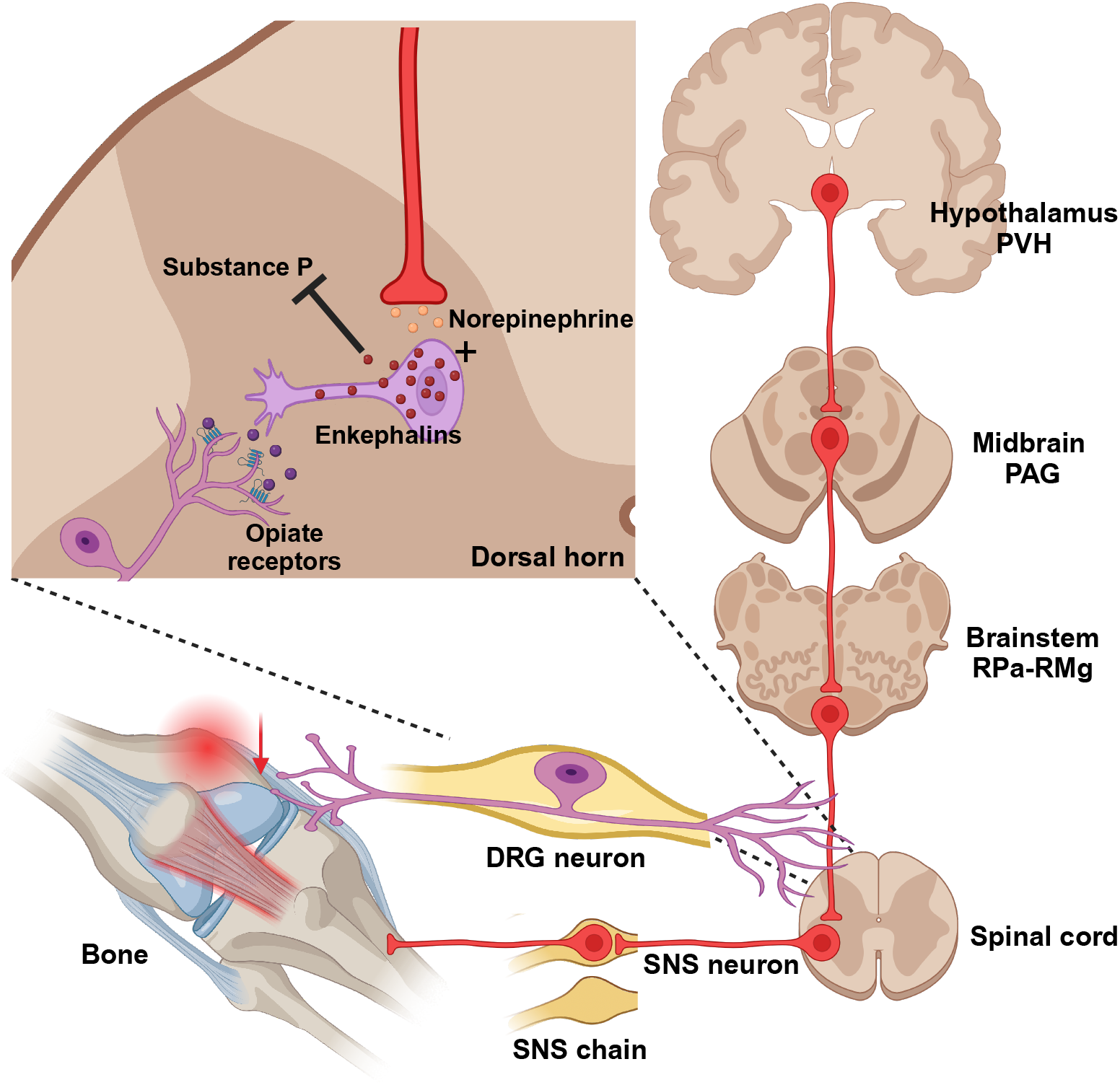
Diagrammatic outline of the SNS brain–bone neuroaxis relevant to pain. The central SNS brain**–**bone circuit starts in the hypothalamic paraventricular nucleus (PVH) known to home SNS pre–autonomic neurons projecting to the SNS neurons of the periaqueductal gray (PAG) in the midbrain. From the PAG the SNS outflow is further relayed to the raphe pallidus— raphe magnus (RPa-RMg) neurons that are terminated in the dorsal horn of spinal gray matter, where they regulate the release of enkephalins that inhibit pain sensation by attenuating substance P (SP) release. In turn, opiates produce antinociception via the *µ*-opiate receptors, in part, through modulation of responses to SP. Neurons in the RMg are involved in the central modulation of noxious stimuli, therefore, the RMg—PAG could be the part of the ascending hierarchical circuit relating to the perception of bone pain.

We also find that two major nuclei in the hypothalamus––the PVH and LH––send SNS efferents to bone. While the PVH, which is a home to major SNS pre–autonomic neurons, sends SNS projections to the bone marrow (Denes *et al*., 2005), we find that LH predominantly innervates the femur. The functions of other hypothalamic SNS–bone feedback circuits are not presently known. However, given that the LH, PVH and DM are main brain regions that send SNS outflow to bone and also express leptin receptors (Flak & Myers, 2016), it is also possible that the anti–osteogenic relay for leptin might originate from neurons in the LH PVH, and/or DM.

Consistent with a prior study (Denes *et al*., 2005), the highest number of PRV152– infected neurons was in the medulla was the RPa and RMg. While we have previously established a contribution of PDE5A–containing neurons in the RPa to bone mass regulation (Kim *et al*., 2020), the functional role of the RMg in regulating bone remains unknown. Whereas projections from the raphe nuclei, including the RPa, terminate in the dorsal horn of spinal gray matter, where they regulate the release of enkephalins that inhibit pain sensation (Francois *et al*, 2017), RMg neurons are involved in the central modulation of noxious stimuli (Fields *et al*, 1991). Thus, the RMg—PAG could be the part of the ascending hierarchical circuit relating to the perception of bone pain. The importance of this circuit for the control of bone pain will require a more comprehensive demonstration of its pervasiveness across and within the mammalian species. Whether or not these findings can be extended to humans, they do provide important actionable targets for pain treatment.

In all, our results provide compelling evidence for a brain—bone SNS neuroaxis, likely part of coordinated and/or multiple redundant mechanisms that regulate bone metabolism and/or nociceptive functions. Furthermore, we show that bone is not innervated by unique neuron groups, but rather by overlapping SNS circuitry common to the control of other peripheral targets, such as bone marrow and adipose tissues. We believe our comprehensive atlas of the brain regions involved in coding and decoding SNS efferent signals to bone would stimulate further research into bone pain and the neural regulation of bone metabolism.

## ACKNOWLEDGEMENTS

Work at Icahn School of Medicine at Mount Sinai carried out at the Center for Translational Medicine and Pharmacology was supported by R01 AG071870 to M.Z., T.Y., and S.-M.K.; R01 AG074092 and U01 AG073148 to T.Y. and M.Z.; and U19 AG060917 and R01 DK113627 to M.Z.

## DATA AVAILABILITY

Figure 2–source data 1 contains the numerical data used to generate the figures.

## Appendix

Glossary of the brain nuclei, sub–nuclei and regions.

### Cerebral cortex

M1: primary motor cortex
M2: secondary motor cortex
S1HL: primary somatosensory cortex, hindlimb region

### Forebrain

AVPe: anteroventral periventricular nucleus
BST: bed nucleus of the stria terminalis
BSTMA: bed nucleus of the stria terminalis, medial division, anterior part
BSTMPL: bed nucleus of stria terminalis, medial division, posterolateral part
BSTMV: bed nucleus of the stria terminalis, medial division, ventral part
LPO: lateral preoptic area
LSI: lateral septal nucleus, intermediate part
LSV: lateral septal nucleus, ventral part
MnPO: median preoptic nucleus
MPOL: medial preoptic nucleus, lateral part
MPOM: medial preoptic nucleus, medial part
SFO: subfornical organ
VMPO: ventromedial preoptic nucleus
VOLT: vascular organ of the lamina terminalis
VP: ventral pallidum

### Thalamus

LHbL: lateral habenular nucleus, lateral part
PrC: precommissural nucleus
pv: periventricular fiber system

### Hypothalamus

AHP: anterior hypothalamic area, posterior part
Arc: arcuate hypothalamic nucleus
ArcLP: arcuate hypothalamic nucleus, lateroposterior part
ArcMP: arcuate hypothalamic nucleus, medial posterior part
BLA: basolateral amygdaloid nucleus, anterior part
CeM: central amygdaloid nucleus, medial division
DM: dorsomedial hypothalamic nucleus
DMD: dorsomedial hypothalamic nucleus, dorsal part
DMV: dorsomedial hypothalamic nucleus, ventral part
LA: lateroanterior hypothalamic nucleus
LH: lateral hypothalamic area
MPA: medial preoptic area
PaAP: paraventricular hypothalamic nucleus, anterior parvicellular part
PaLM: paraventricular hypothalamic nucleus, lateral magnocellular part
PaMM: paraventricular hypothalamic nucleus, medial magnocellular part
PaMP: paraventricular hypothalamic nucleus, medial parvicellular part
PaPo: paraventricular hypothalamic nucleus, posterior part
PaV: paraventricular hypothalamic nucleus, ventral part
PH: posterior hypothalamic area
PVH: paraventricular hypothalamic nucleus
SCh: suprachiasmatic nucleus
SChVL: suprachiasmatic nucleus, ventrolateral part
TC: tuber cinereum area
VMHC: ventromedial hypothalamic nucleus, central part
ZI: zona incerta

### Midbrain and pons

CGPn: central gray of the pons
CnF: cuneiform nucleus
DLPAG: dorsolateral periaqueductal gray
DMPAG: dorsomedial periaqueductal gray
DMTg: dorsomedial tegmental area
DpMe: deep mesencephalic nucleus
DRD: dorsal raphe nucleus, dorsal part
DRI: dorsal raphe nucleus, interfascicular part
LC: locus coeruleus
LDTg: laterodorsal tegmental nucleus
LPAG: lateral periaqueductal gray
Mo5: motor trigeminal nucleus
MPB: medial parabrachial nucleus
PAG: periaqueductal gray
PnO: pontine reticular nucleus, oral part
SubCV: subcoeruleus nucleus, ventral part
VLPAG: ventrolateral periaqueductal gray
VTA: ventral tegmental area

### Medulla

7N: facial nucleus
DPGi: dorsal paragigantocellular nucleus
Gi: gigantocellular reticular nucleus
GiA: gigantocellular reticular nucleus, alpha part
GiV: gigantocellular reticular nucleus, ventral part
IRt: intermediate reticular nucleus
LPGi: lateral paragigantocellular nucleus
MVePC: medial vestibular nucleus, parvicellular part
NTS: nucleus of the solitary tract
Pr: prepositus nucleus
RMg: raphe magnus nucleus
ROb: raphe obscurus nucleus
RPa: raphe pallidus nucleus
SolC: nucleus of the solitary tract, commissural part
SolCe: nucleus of the solitary tract, central part
SolDL: solitary nucleus, dorsolateral part
SolDM: nucleus of the solitary tract, dorsomedial part
SolG: nucleus of the solitary tract, gelatinous part
SolI: nucleus of the solitary tract, interstitial part
SolIM: nucleus of the solitary tract, intermediate part
SolM: nucleus of the solitary tract, medial part
SolV: solitary nucleus, ventral part
SolVL: nucleus of the solitary tract, ventrolateral part

## REFERENCES

Adler ES, Hollis JH, Clarke IJ, Grattan DR, Oldfield BJ (2012) Neurochemical characterization and sexual dimorphism of projections from the brain to abdominal and subcutaneous white adipose tissue in the rat. J Neurosci 32: 15913–15921

Bamshad M, Aoki VT, Adkison MG, Warren WS, Bartness TJ (1998) Central nervous system origins of the sympathetic nervous system outflow to white adipose tissue. Am J Physiol 275: R291–299

Bamshad M, Song CK, Bartness TJ (1999) CNS origins of the sympathetic nervous system outflow to brown adipose tissue. Am J Physiol 276: R1569–1578

Baptista-de-Souza D, Pelarin V, Canto-de-Souza L, Nunes-de-Souza RL, Canto-de-Souza A (2018) Interplay between 5-HT(2C) and 5-HT(1A) receptors in the dorsal periaqueductal gray in the modulation of fear-induced antinociception in mice. Neuropharmacology 140: 100–106

Benarroch EE (2008) Descending monoaminergic pain modulation: bidirectional control and clinical relevance. Neurology 71: 217–221

Bliziotes M, McLoughlin S, Gunness M, Fumagalli F, Jones SR, Caron MG (2000) Bone histomorphometric and biomechanical abnormalities in mice homozygous for deletion of the dopamine transporter gene. Bone 26: 15–19

Bowers RR, Festuccia WT, Song CK, Shi H, Migliorini RH, Bartness TJ (2004) Sympathetic innervation of white adipose tissue and its regulation of fat cell number. Am J Physiol Regul Integr Comp Physiol 286: R1167–1175

Calvino B, Grilo RM (2006) Central pain control. Joint Bone Spine 73: 10–16

Curanovic D, Enquist L (2009) Directional transneuronal spread of alpha-herpesvirus infection. Future Virol 4: 591

Denes A, Boldogkoi Z, Uhereczky G, Hornyak A, Rusvai M, Palkovits M, Kovacs KJ (2005) Central autonomic control of the bone marrow: multisynaptic tract tracing by recombinant pseudorabies virus. Neuroscience 134: 947–963

Ducy P, Amling M, Takeda S, Priemel M, Schilling AF, Beil FT, Shen J, Vinson C, Rueger JM, Karsenty G (2000) Leptin inhibits bone formation through a hypothalamic relay: a central control of bone mass. Cell 100: 197–207

Ekstrand MI, Enquist LW, Pomeranz LE (2008) The alpha-herpesviruses: molecular pathfinders in nervous system circuits. Trends Mol Med 14: 134–140

Elefteriou F (2018) Impact of the Autonomic Nervous System on the Skeleton. Physiol Rev 98: 1083–1112

Elefteriou F, Ahn JD, Takeda S, Starbuck M, Yang X, Liu X, Kondo H, Richards WG, Bannon TW, Noda M, Clement K, Vaisse C, Karsenty G (2005) Leptin regulation of bone resorption by the sympathetic nervous system and CART. Nature 434: 514–520

Enquist LW (2002) Exploiting circuit-specific spread of pseudorabies virus in the central nervous system: insights to pathogenesis and circuit tracers. J Infect Dis 186 Suppl 2: S209–214

Fields HL, Heinricher MM, Mason P (1991) Neurotransmitters in nociceptive modulatory circuits. Annu Rev Neurosci 14: 219–245

Flak JN, Myers MG, Jr. (2016) Minireview: CNS Mechanisms of Leptin Action. Mol Endocrinol 30: 3–12

Francis NJ, Asmus SE, Landis SC (1997) CNTF and LIF are not required for the target-directed acquisition of cholinergic and peptidergic properties by sympathetic neurons in vivo. Dev Biol 182: 76–87

Francois A, Low SA, Sypek EI, Christensen AJ, Sotoudeh C, Beier KT, Ramakrishnan C, Ritola KD, Sharif-Naeini R, Deisseroth K, Delp SL, Malenka RC, Luo L, Hantman AW, Scherrer G (2017) A Brainstem-Spinal Cord Inhibitory Circuit for Mechanical Pain Modulation by GABA and Enkephalins. Neuron 93: 822–839 e826

Hill EL, Elde R (1991) Distribution of CGRP-, VIP-, D beta H-, SP-, and NPY-immunoreactive nerves in the periosteum of the rat. Cell Tissue Res 264: 469–480

Hohmann EL, Elde RP, Rysavy JA, Einzig S, Gebhard RL (1986) Innervation of periosteum and bone by sympathetic vasoactive intestinal peptide-containing nerve fibers. Science 232: 868–871

Kim SM, Taneja C, Perez-Pena H, Ryu V, Gumerova A, Li W, Ahmad N, Zhu LL, Liu P, Mathew M, Korkmaz F, Gera S, Sant D, Hadelia E, Ievleva K, Kuo TC, Miyashita H, Liu L, Tourkova I, Stanley S, et al (2020) Repurposing erectile dysfunction drugs tadalafil and vardenafil to increase bone mass. Proc Natl Acad Sci U S A 117: 14386–14394

Leitner C, Bartness TJ (2009) Acute brown adipose tissue temperature response to cold in monosodium glutamate-treated Siberian hamsters. Brain Res 1292: 38–51

Mantyh PW (1982) The ascending input to the midbrain periaqueductal gray of the primate. J Comp Neurol 211: 50–64

Martin CD, Jimenez-Andrade JM, Ghilardi JR, Mantyh PW (2007) Organization of a unique net-like meshwork of CGRP+ sensory fibers in the mouse periosteum: implications for the generation and maintenance of bone fracture pain. Neurosci Lett 427: 148–152

Nguyen NL, Randall J, Banfield BW, Bartness TJ (2014) Central sympathetic innervations to visceral and subcutaneous white adipose tissue. Am J Physiol Regul Integr Comp Physiol 306: R375–386

Pechura CM, Liu RP (1986) Spinal neurons which project to the periaqueductal gray and the medullary reticular formation via axon collaterals: a double-label fluorescence study in the rat. Brain Res 374: 357–361

Ryu V, Bartness TJ (2014) Short and long sympathetic-sensory feedback loops in white fat. Am J Physiol Regul Integr Comp Physiol 306: R886–900

Ryu V, Garretson JT, Liu Y, Vaughan CH, Bartness TJ (2015) Brown adipose tissue has sympathetic-sensory feedback circuits. J Neurosci 35: 2181–2190

Ryu V, Watts AG, Xue B, Bartness TJ (2017) Bidirectional crosstalk between the sensory and sympathetic motor systems innervating brown and white adipose tissue in male Siberian hamsters. Am J Physiol Regul Integr Comp Physiol 312: R324–R337

Shi H, Bartness TJ (2001) Neurochemical phenotype of sympathetic nervous system outflow from brain to white fat. Brain Res Bull 54: 375–385

Song CK, Bartness TJ (2001) CNS sympathetic outflow neurons to white fat that express MEL receptors may mediate seasonal adiposity. Am J Physiol Regul Integr Comp Physiol 281: R666–672

Song CK, Enquist LW, Bartness TJ (2005a) New developments in tracing neural circuits with herpesviruses. Virus Res 111: 235–249

Song CK, Jackson RM, Harris RB, Richard D, Bartness TJ (2005b) Melanocortin-4 receptor mRNA is expressed in sympathetic nervous system outflow neurons to white adipose tissue. Am J Physiol Regul Integr Comp Physiol 289: R1467–1476

Song CK, Vaughan CH, Keen-Rhinehart E, Harris RB, Richard D, Bartness TJ (2008) Melanocortin-4 receptor mRNA expressed in sympathetic outflow neurons to brown adipose tissue: neuroanatomical and functional evidence. Am J Physiol Regul Integr Comp Physiol 295: R417–428

Takeda S, Elefteriou F, Levasseur R, Liu X, Zhao L, Parker KL, Armstrong D, Ducy P, Karsenty G (2002) Leptin regulates bone formation via the sympathetic nervous system. Cell 111: 305–317

Vaughan CH, Bartness TJ (2012) Anterograde transneuronal viral tract tracing reveals central sensory circuits from brown fat and sensory denervation alters its thermogenic responses. Am J Physiol Regul Integr Comp Physiol 302: R1049–1058

